# 3D melanoma spheroid model for the development of positronium biomarker

**DOI:** 10.1101/2022.06.01.494339

**Authors:** Hanieh Karimi, Paweł Moskal, Ewa Ł. Stępień

## Abstract

It was recently demonstrated that newly invented positronium imaging may be used for improving cancer diagnostics by providing additional information about tissue pathology with respect to the standardized uptake value currently available in positron emission tomography (PET). Positronium imaging utilizes properties of a positronium atoms, which are built from the electron and positron produced in the body during PET examinations.

We hypothesized whether positronium imaging would be sensitive to *in vitro* discrimination of tumour-like three-dimensional structures (spheroids) build of melanoma cell lines with different cancer activity and biological properties.

The lifetime of ortho-Positronium (o-Ps) was evaluated in melanoma spheroids from two cell lines (WM266-4 and WM115) differing in the stage of malignancy. Additionally, we considered such parameters: as cell size, proliferation rate and malignancy to evaluate their relationship with o-Ps lifetime. We demonstrate the pilot results for the o-Ps lifetime measurement in extracellular matrix free spheroids. With the statistical significance of two standard deviations, we demonstrated that the higher the degree of malignancy and the rate of proliferation of neoplastic cells the shorter the lifetime of ortho-positronium. In particular we observed following indications encouraging further research: (i) WM266-4 spheroids characterized with higher proliferation rate and malignancy showed shorter o-Ps lifetime compared to WM115 spheroids characterized by lower growth rate, (ii) Both cell lines showed a decrease in the lifetime of o-Ps after spheroid generation in 8^th^ day comparing to 4^th^ day in culture and the mean o-Ps lifetime is longer for spheroids formed from WM115 cells than these from WM266-4 cells, regardless spheroid age. The results of these study revealed that positronium is a promising biomarker that may be applied in PET diagnostics for the assessment of the degree of cancer malignancy.

## Introduction

Over the past few decades three-dimensional (3D) cell cultures have been widely used as *in vitro* models which can bridge the gap between *in vitro* and *in vivo* cell conditions [1]. The comparison of the 3D cell culture to a cell monolayer revealed some specific physiological and morphological characteristics such as cell-to-cell, and cell-matrix interactions, cell signaling, proliferation and necrosis. Unlike a monolayer cell culture, 3D spheroid is an appropriate model to mimic the real tumor cells environment and nutrient diffusion rate between cells. Multicellular tumor spheroids (MCTS) allow to study the biochemical mechanism of cell growth, enzymatic reactions, and various treatment modalities [2-4].

Melanoma is a prevalent type of skin cancer that has been categorized as one of the most lethal cancers. Melanoma develops from melanocytes – pigmented cells producing melanin and, located in an epidermis-due to their genetic modifications [5], (Fig 1).

**Figure 1.**
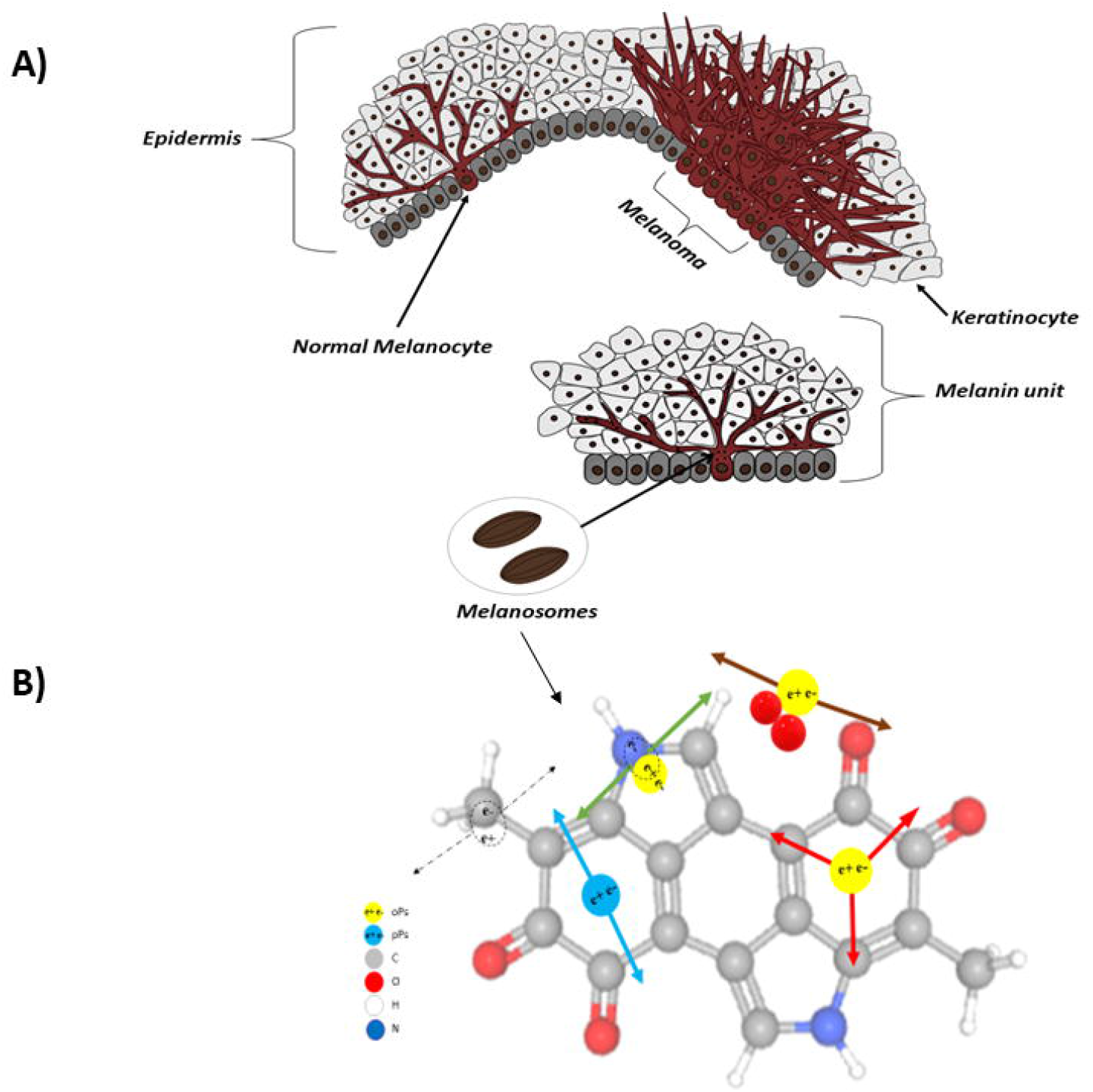
**A) Schematic view of melanocyte cells and the formation of melanoma in epidermis layer**. Normal melanocyte cells have regular shape and grow in a controlled way but after modification to cancer cells, they grow on the top of each other, invade to neighbour cells and have irregular and uncontrolled division. **B) Pictorial illustrations of positron annihilations in the melanin molecules**. Carbon, Oxygen, Hydrogen, Nitrogen, p-Ps and o-Ps atoms are indicated in colours explained in the legend. Dashed arrows indicate photons from direct electron-positron annihilation. Red arrows indicate photons from o-Ps atom self-annihilation. Blue arrows show photons from p-Ps self-annihilation. Green and brown arrows indicate o-Ps decay via pick-off and conversion processes, respectively.

Although numerous studies have been dedicated to diagnosing melanoma in the early stages [6], it still needs more investigation and follow up. The rate of metastasis in patients due to melanoma is high, 350,000 new cases per year, and the number of new cases is significantly increasing each year [7–9].

In this study, 3D spheroid model of two melanoma cell lines, WM266-4 and WM115, with different stage of malignancy, have been evaluated. Melanoma cell lines WM266-4 and WM115 have various biological properties which make them different in proliferation, migration rate, and cell size. The activation of advanced glycation end products (AGEs) which are produced during combination of fat and protein with sugar in human body and proteins such as receptor for advanced glycation end products (RAGE), and c-Jun N-terminal kinases (JNK) in the WM266-4 line makes this cell line more invasive in comparison to the WM115 line [10]. The WM115 cell line has higher pigmentation which makes it darker in comparison to the WM266-4 cell line

In this study we hypothesized whether the difference between the grade of malignancy of the WM115 and WM266-4 melanoma cancers, present at the level of the cell physiology can be probed by positronium biomarker. Positronium is an exotic atom built of positron and electron. Positronium is formed also in the intra-molecular spaces during the PET diagnosis [11]. In the tissue, positronium may be formed and trapped in the free voids of the intra-molecular spaces, as it is shown pictorially in (Fig 1). Positron emitted from the radionuclide (eg. ^18^F in the PET diagnosis or ^22^Na in the typical Positron Annihilation Lifetime Spectroscopy (PALS) experiments) penetrates the object, and after losing the energy annihilates with electron from the molecules constituting the cells. Positron electron annihilation into photons may occur directly (e+e- -> photons (black dashed arrows in (Fig 1))) or *via* positronium atom (e+e- -> positronium -> photons (solid arrows in (Fig 1))). In quarter of cases positronium is formed as short-lived (125 ps) state called para-positronium (p-Ps) and in three quarter of cases as long-lived (142 ns) ortho-positronium (o-Ps). Para-positronium (indicated in blue in (Fig 1)) decays predominantly into two photons (blue arrows) and ortho-positronium decays in vacuum predominantly into three photons (red arrows). However, in the intra-molecular voids ortho-positronium undergoes processes as pick-off (positron from o-Ps annihilates into two photons (green arrows) with electron from the surrounding molecule) and conversion into para-positronium *via* interaction with molecules as e.g., oxygen molecule. The resulting para-positronium decays into two photons (brown arrows). The range of the o-Ps mean lifetime variation is significant, and it changes from the value of 142 ns in vacuum to 1.8 ns in water [12, 13]. Therefore, ortho-positronium lifetime depends strongly on the molecular environment: the nano-molecular structure and concentration of the bio-active molecules [11-18]. Properties of positronium in biological samples are only scarcely studied so far. Only recently, the first studies of positronium in the 3D cell structures were performed by culturing cell on collagen matrix [19]. As regards skin cells, they were investigated with low energy positron beam [20-22].

The first in-vitro research for the positronium lifetime in the tissues operated from the patients indicate promising results in view of application of positronium as a biomarker for the diagnosis of uterine cancer and myxoma cancer and as a biomarker of hypoxia [17,18, 23,24]. Recently positronium was proposed as a novel biomarker for the in-vivo assessment of tissue pathology [11, 24] that can be imaged using newly developed positronium imaging method [12,16,17,25] when applying prompt photon radionuclides [26-28] and high sensitivity PET scanners [29]. Moreover, the advent of total-body PET systems characterized by high sensitivity [29-32] will enable to apply positronium imaging simultaneously to the standard metabolic imaging during the Positron Emission Tomography [29].

The aim of this study was to check whether positronium imaging would be sensitive to *in vitro* discrimination of tumour-like three-dimensional structures (spheroids) build of melanoma cell lines with different cancer activity and biological properties.

## Results

### Spheroid morphology and proliferation rate

The exemplary spheroids grown in the micro plate are shown in (Fig 2). WM266-4 cell lines formed more spherical and concentrated spheroids in comparison to WM115 cell line. Spheroids in both cell lines showed an increase in their size and circularity over the time. Generally, the rate of growth and size of spheroids depend on the size of cavities and the type of plates that have been seeded. The bigger micro cavities, the bigger spheroids will be formed. Although the size of spheroids can be controlled by initial culture seeding, the scales of micro wells can also affect the spheroid diameter [33].

**Figure 2.**
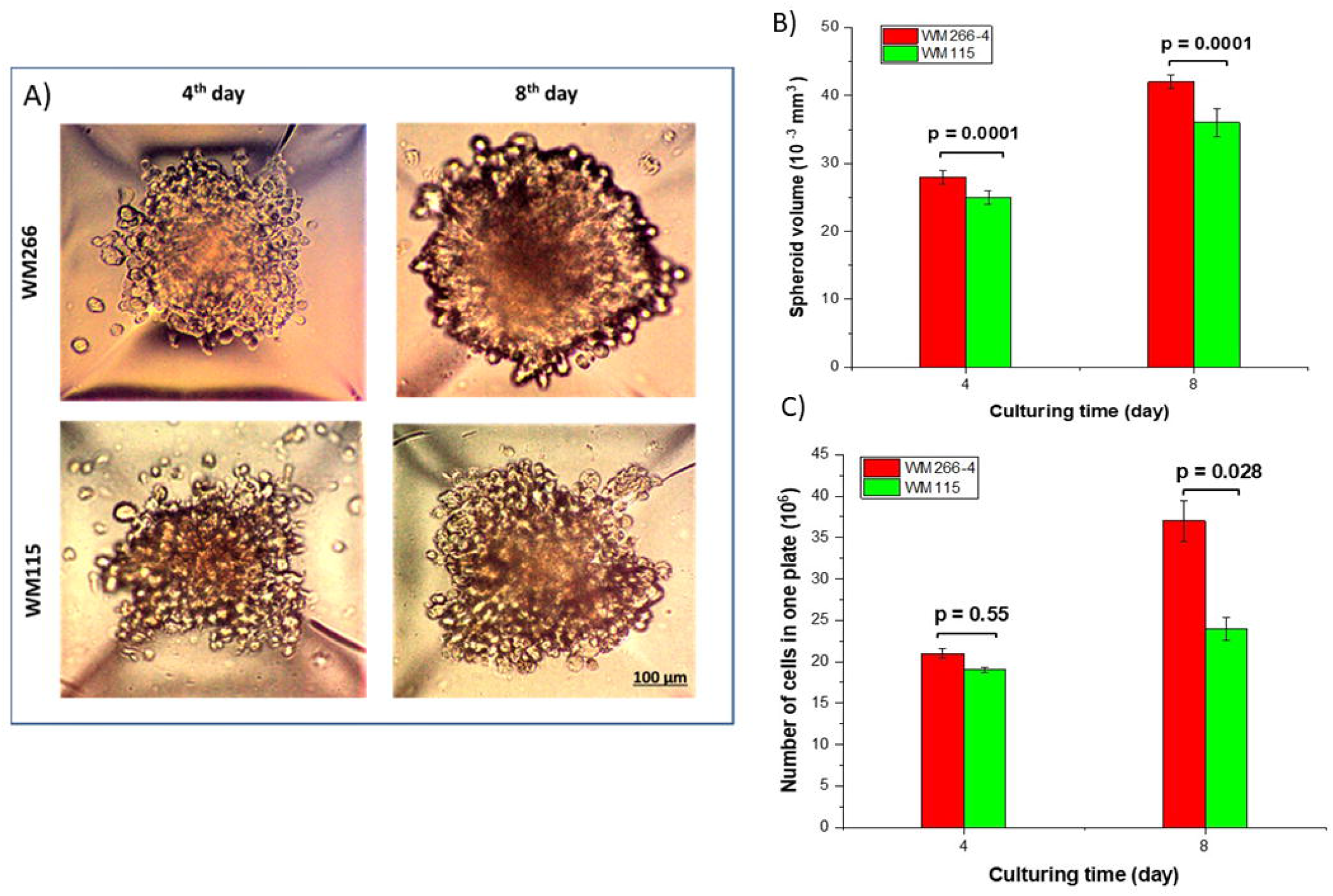
A) Microscopic images of two different cell lines, WM266-4 and WM115, the 4^th^ and 8^th^ day of culturing. The density and circularity of spheroids is increasing during the culturing time. B) Spheroids volume as a function of culturing time: Spheroids have grown during the time. WM266-4 has a faster proliferation due to its higher malignant level than WM115. C) Number of cells in plate as the function of culturing time: The number of cells has increased during the culturing time and WM266-4 as malignant 3D cell culture shows a higher velocity in proliferation and division. The number of cells in one plate has been calculated using the Luna II cell counter machine after spheroid harvesting.

Figure 2. (B, C) shows that WM266-4 and WM115 spheroids have grown during the time. WM266-4 spheroids exhibited a faster proliferation rate than WM115 which backs to its malignant characteristics.

Figure 2. (B) presents, that there is an increase in the number of cells from cell seeding until the 8^th^ day after culturing. The division rate of WM266-4 cells in the spheroid is higher than these of WM115. For WM266-4, it results in 1.5 and 2.74-fold increase of cell number after 4^th^ and 8^th^ day, respectively. Number of cells in WM115 spheroids increased 1.4 and 1.7-fold after 4 and 8 days, respectively. In each positronium lifetime measurement, a total of 36000 spheroids were used.

### Spheroids’ cells characterisation

WM266-4 spheroids comprise cells with diameter of 15.70 ± 0.10 μm at day 4 after cell seeding and 15.92 ± 0.08 μm at day 8, while WM155 spheroid cells diameter was equal to 16.66 ± 0.20 μm at day 4 and 17.28 ± 0.25 μm at day 8. The mean size of stained cells in 2D cell culture was smaller than size of cells from the 3D spheroids, 14.65 ± 0.09 μm and 16.27 ± 0.10 μm for WM266-4 and WM115, respectively.

The number of cells in both cell lines were growing in time. The observed growth was faster for WM266-4 cell line than for the WM115. The WM115 has longer doubling time than WM266-4, approximately 7.5 and 6 days, respectively, which means cells of WM115 line spend more time in their cell cycle with respect to cells from WM266-4 line. The doubling time (DT) of spheroids is usually characterized like real tumour doubling time and is calculated by using spheroid volume *via*:

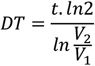

Where V_1_ and V_2_ are spheroid volumes at times t_1_ and t_2_ = t_1_+ t after seeding, respectively [34-36].

### The lifetime of ortho-positronium

The mean lifetime of ortho-positronium atoms in the spheroids was established for two different cell lines characterized with different degree of malignancy. Both measurements were performed for two different ages of spheroids: at fourth and eight days after seeding. The viability tests have been evaluated before and after each PALS measurement. The tests showed that spheroids viability remained constant above 85% till the 8^th^ day. Thus, spheroids were in good and stable condition during the experiment. Each viability test has been done on 4 different samples in each measurement under the same condition.

The measurement of positronium lifetime have been repeated three times. In each measurement, spheroids have been harvested from the plate and put inside the chamber.

Figure 3. demonstrates the results of mean o-Ps lifetime and distribution in 3D melanoma spheroids with different malignancy levels at two different ages.

**Figure 3.**
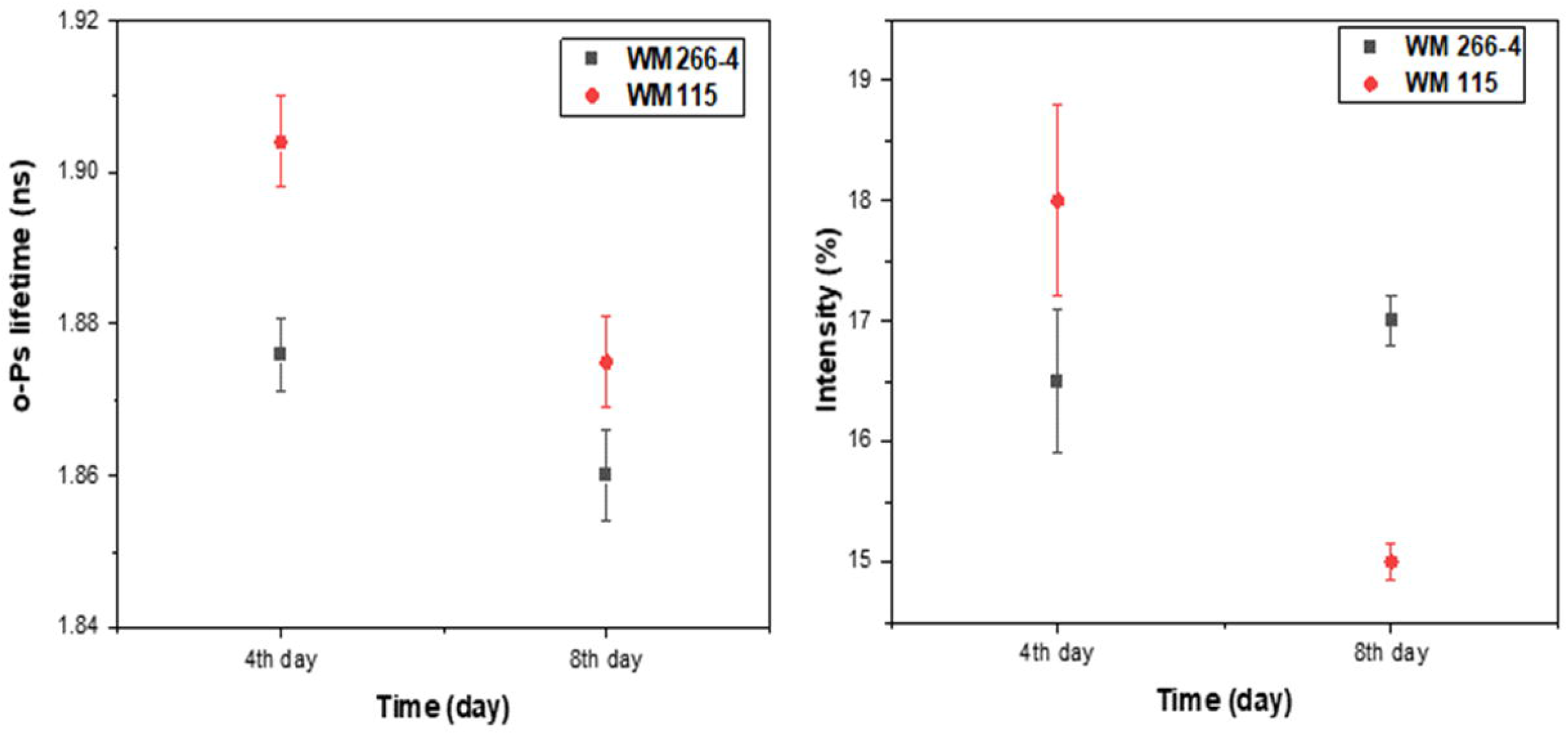
(Left) The lifetime of o-Ps in WM266-4 (black squares) and WM115 (red dots) spheroids in two different ages, 4 and 8 days after cell seeding. (Right) Intensity of o-Ps production in WM266-4 spheroids (black squares) and WM115 spheroids (red dots) as a function of time.

The obtained results given in (Table 1) show that WM266-4 spheroids with more malignancy, proliferation rate, and high concentration of cells in spheroids have a shorter ortho-positronium mean lifetime than WM115 cells, therefore, smaller free inter-molecular voids, than WM115 cells from the primary tumor with less division rate and lower concentration of cells over time. Although both cell lines show a gradual decrease in the lifetime of o-Ps during the time of culturing, o-Ps shows a longer lifetime in WM115 than in WM266-4 spheroids. The Intensity in WM266-4 spheroids remained almost constant while WM115 spheroids present a decrease in the intensity which can be considered as changes at the molecular level in WM115 spheroids cells. The lifetimes of o-Ps in the cell can be different from the lifetime measured in the case when the cell is in the medium containing water, collagen, or other chemical compounds because the cell surface, cell growth, and the shape of cells can be changed. Therefore, in this research, only spheroids without medium, water, and any chemicals have been evaluated.

**Table 1.** o-Ps mean lifetime (τ_o-Ps_) and production intensity (I) in 3D melanoma spheroids without medium and chemical compounds.

| Cell line | 4<sup>th</sup> day | 8<sup>th</sup> day |
| --- | --- | --- |
| | $\tau_{o-Ps}$ [ns], I (%) | $\tau_{o-Ps}$ [ns], I (%) |
| WM266-4 | 1.876 (0.005), 16.5% (0.6) | 1.861 (0.006), 17.0% (0.2) |
| WM115 | 1.909 (0.006), 18.0% (0.8) | 1.875 (0.006), 15.0% (0.2) |

## Methods

### Cell culture

Two melanoma cell lines, WM266-4, a human malignant melanoma cancer cell line and WM115, a cell line from primary skin cancer were purchased from ESTDAB Melanoma Cell Bank (Tübingen, Germany) and cultured as we previously described [37]. Luna-II™ automated cell counter (Logos Biosystems, Inc.) was used to determine the cell counts and viability before cell seeding.

### Spheroid generation

Both cell lines, WM266-4 and WM115, have been seeded in 5D spherical plates (5D sp5dplate, Kugelmeiers, Switzerland) to form spheroids. The 5D microplate has 24 wells, 12 wells for the spheroid formation, and 12 wells as control. Each well contains 750 microcavities that are separated from each other by sharp borders and these borders prevent cell migration from one microcavity to another one, thus 9000 spheroids with uniform shape and diameter can be cultured in a single plate.

For cell seeding, firstly 0.5 ml complete medium was added to each well. Then extra 0.5 ml medium including 1 125 000 cells/well (1500 cells/microwell) was added to each well of the plate. The cells in the falcon were resuspended to distribute them in the whole medium and then added to the wells, as it is presented pictorially in (Fig 4).

**Figure 4.**
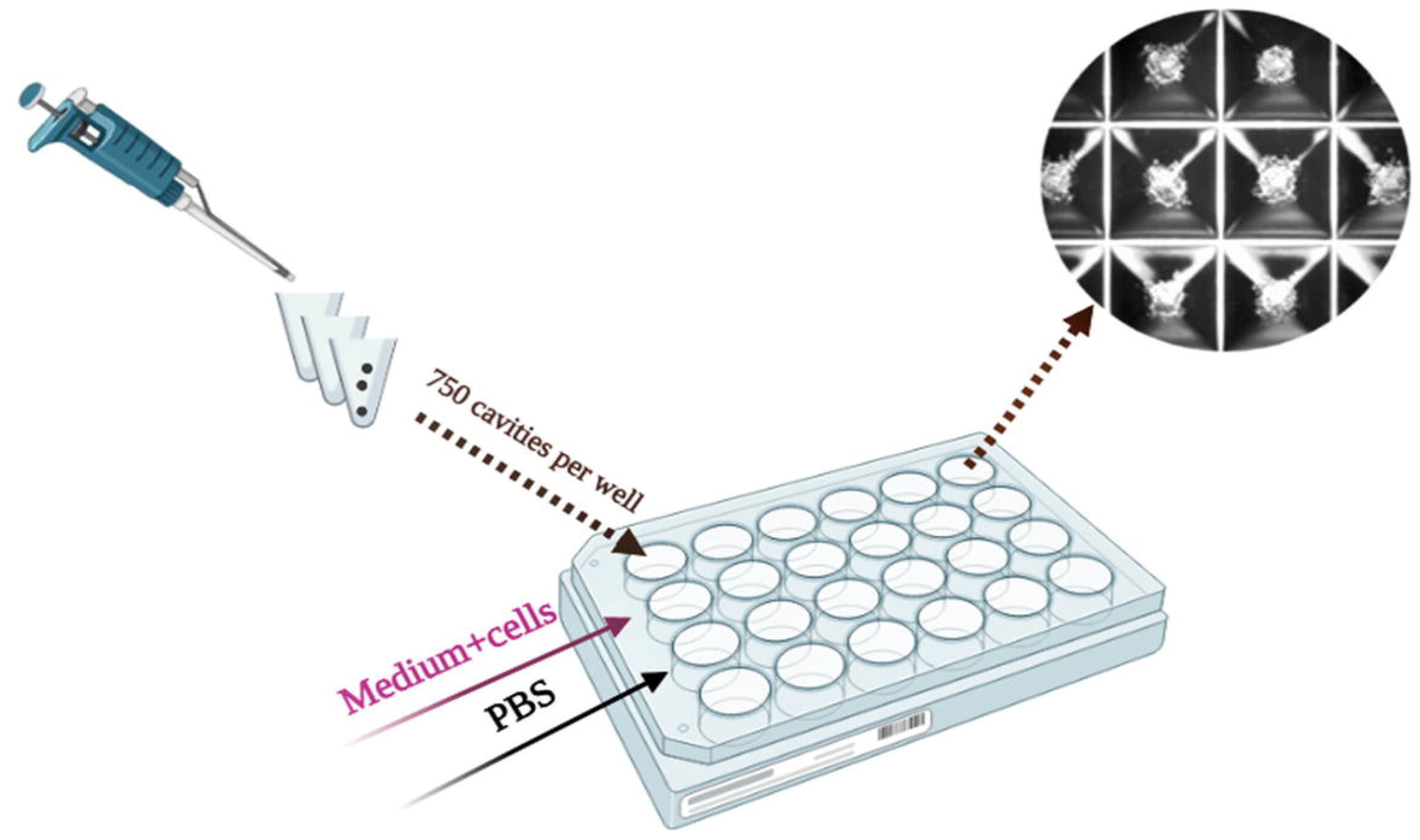
Spheroid formation using 5D microplate. Spheroids were formed 2 to 3 days after cell seeding depending on the cell line. The upper right picture shows spheroids inside part of microcavities (with 509 μm diameter and 320 μm depth).

After cell seeding, plates were maintained in an incubator at 37 °C and 5% CO_2_ with humidity. Cell culture medium was renewed every day after the spheroids were formed. Two suitable time points of spheroid growth were chosen to measure the mean o-Ps lifetime. Spheroid’s morphology and proliferation rate were also determined under the optical microscope (Olympus, IX-LWPO, T2, Japan) at 4^th^ and 8^th^ days after cell seeding. Image analysis was conducted by ImageJ software.

### Viability test assay

Firstly, spheroids were dissociated to single-cell suspension with trypsin/EDTA (cat. no. 25200072, Waltham, MA USA). 100μl of trypsin/EDTA Spheroids were incubated for 10-20 minutes at 37°C and pipetted several times to separate the cells. Then cells were centrifuged at 300 g for 3 min, the supernatant was removed, 100μl medium (Gibco, cat. no. 10010056, Waltham, MA, USA) was added to cells and pipetted several times to separate cells completely. In the final step, 10μl of cells were added to 10 μl trypan blue and counted (Luna-II™ cell counter, logos biosystems). The workflow is shown in (Fig 5).

**Figure 5.**
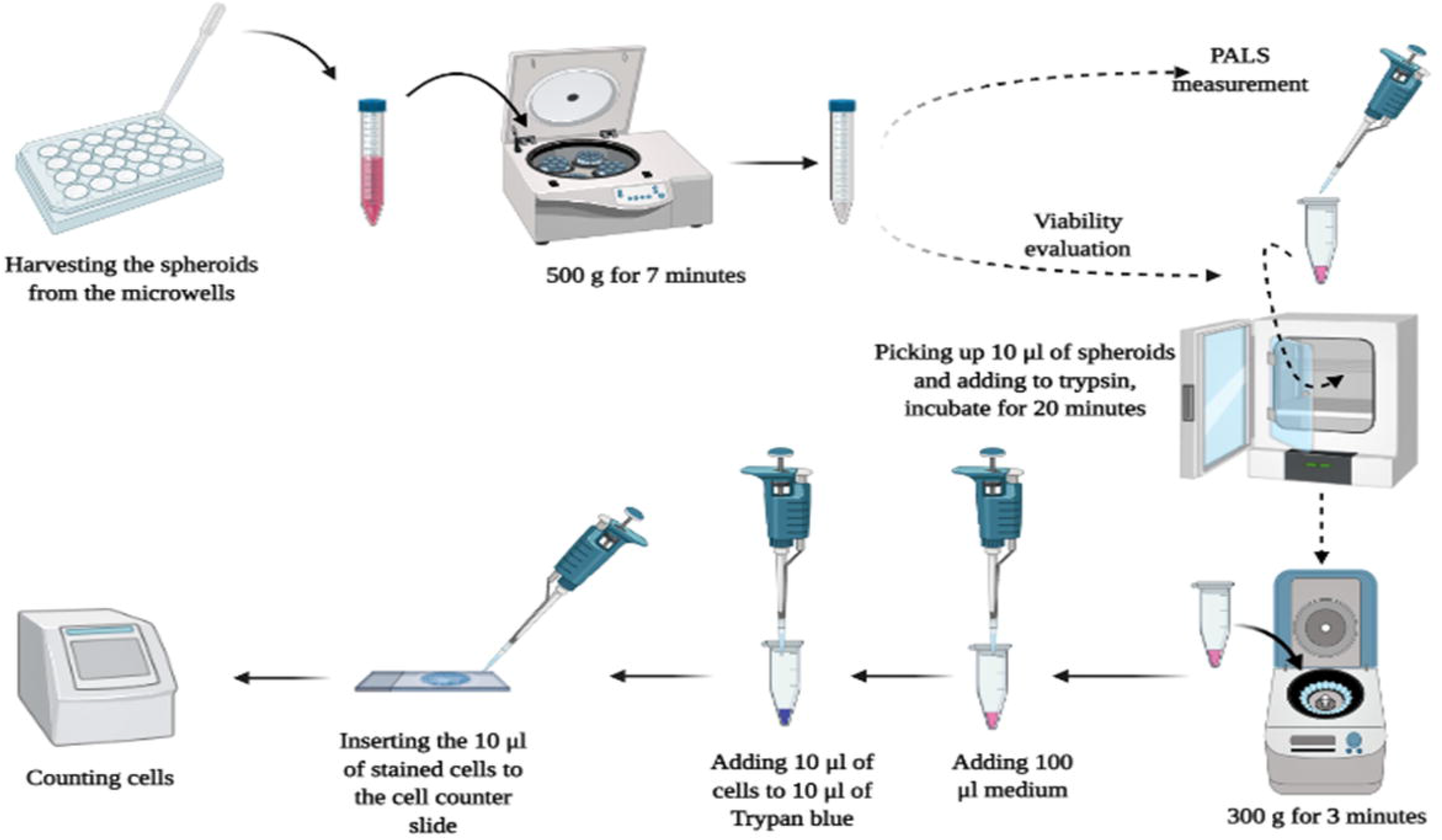
Workflow for investigation of spheroid viability after harvesting 3D spheroids from 5D microplates.

#### Estimation of cell size

After 15 minutes in the incubator, the cells were checked to ensure there were no aggregates. After the viability test using the cell counter, the cell cluster map has been evaluated to see the percentage of clusters. In our experiments, cluster maps showed a high percentage of dissociation, above 90%. Afterward, the size of cells for both cell lines in 3D shape has been checked by cell counter and microscopic investigations. For estimation of the cell size, Luna II ™ cell counter has been used. Regarding the histogram of size distribution, the mean diameter of cells was estimated. For the comparison, the size of cells in 2D cultures was also determined. For this purpose, spheroids were removed from each well using a 3 ml pipette, with a large bore to not destroy the spheroids’ structure. Then spheroids were poured onto a 15ml Falcon (cat. no. 601052NEST/G66010522, GenoPlast, Biochemicals) and centrifuged in 500 g for 7 minutes. In the next step, the supernatant was removed, and 1ml fresh medium added to spheroids.

### Positron annihilation lifetime spectroscopy in 3D spheroids

For obtaining reliable and precise results of positronium lifetime in spheroids, we used spheroids without any medium, supernatant, and chemical compounds. In this method, harvested spheroids from the microplates were prepared for positronium lifetime measurement after centrifuge and removing the medium completely from spheroids.

Positronium lifetime was measured using spectrometer consisting of two vertically arranged BaF_2_ plastic scintillators (SCIONIX, Holland) and two photomultipliers with serial numbers SBO696 and SBO697 (Hamamatsu, Japan) powered by the high voltage power supply (CAEN SY4527). The signals from photomultipliers were read out and analysed by the 6000A data analyzer oscilloscope (Le Croy). Spheroids were irradiated by positrons emitted from the ^22^Na radionuclide (with activity of about 1 MBq) placed in Kapton foil. A dedicated aluminum chamber was used as a container for spheroids and a holder which was connected to a heater to keep cells at 37^0^C. The measurement setup is shown pictorially in (Fig 6).

**Figure 6.**
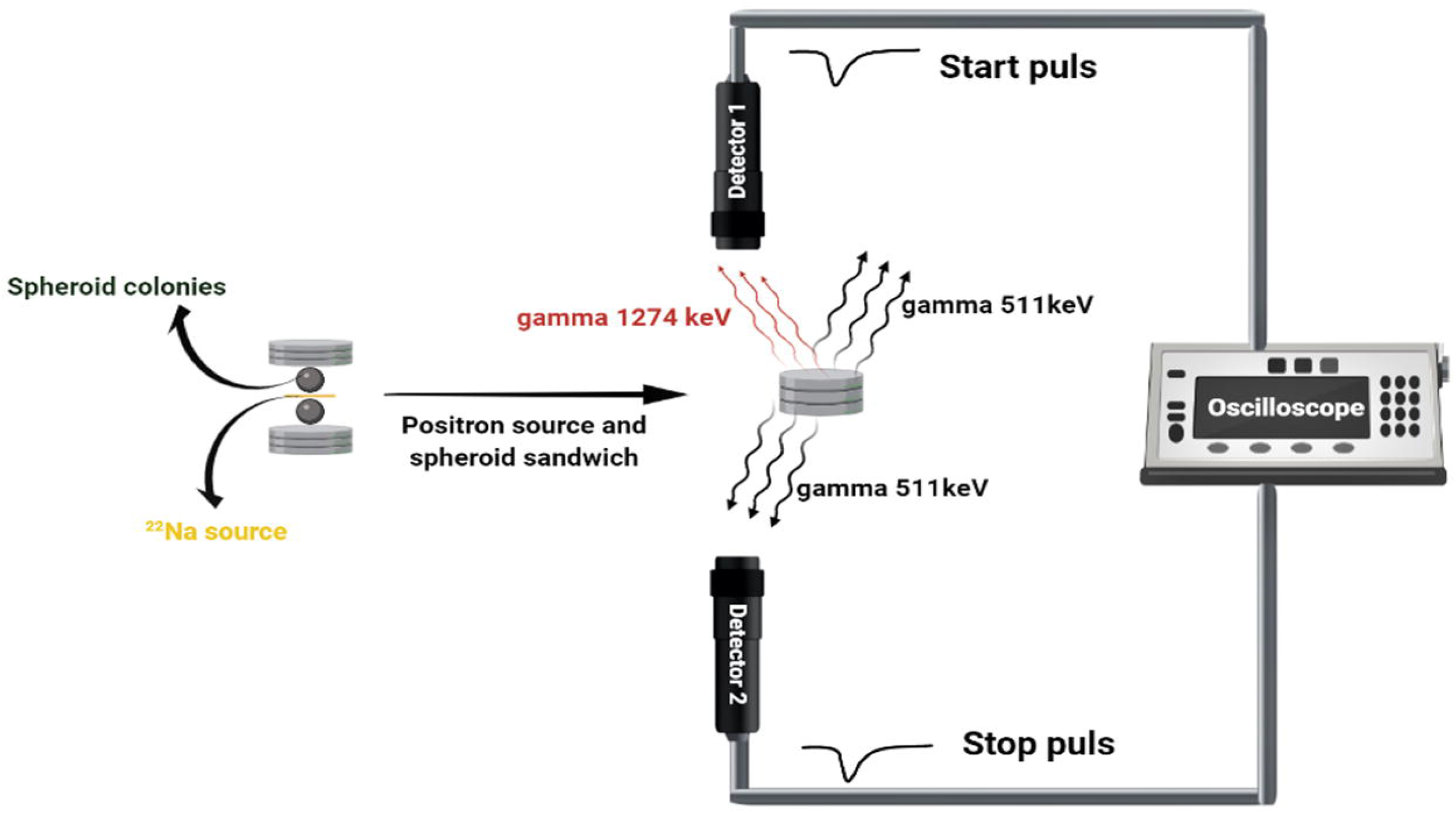
Spheroids surrounding the ^22^Na source are located in the dedicated chamber between two BaF_2_ detectors. Spheroids are adjacent to the ^22^Na source and there is no space or bubble between them. For each measurement 10^6^ events with the coincident registration of 511 KeV photon and 1274 keV photon were collected.

^22^Na radionuclide after emission of positron transforms to the excited state of the ^22^Ne isotope that deexcites (on the average after 2.6 ps) *via* emission of the 1274 KeV gamma quantum [26,27]. Positron loses energy while passing through the cells and eventually annihilates with the electron into two back-to-back 511 keV gamma quanta. The positron-electron annihilation may proceed directly or *via* creation of the positronium. The time between the emission of positron and its annihilation is measured by the registration of the the 1274 KeV deexcitation photon and one of the 511 keV annihilation photons. The sample with the source is positioned in a way (see Fig 6) that enables coincident registration of 1274 keV gamma and one 511 keV photon, and it prevents from the coincident registration of both 511 keV photons flying back-to-back. An exemplary lifetime spectrum determined as a result of the measurement is shown in (Fig 7).

**Figure 7.**
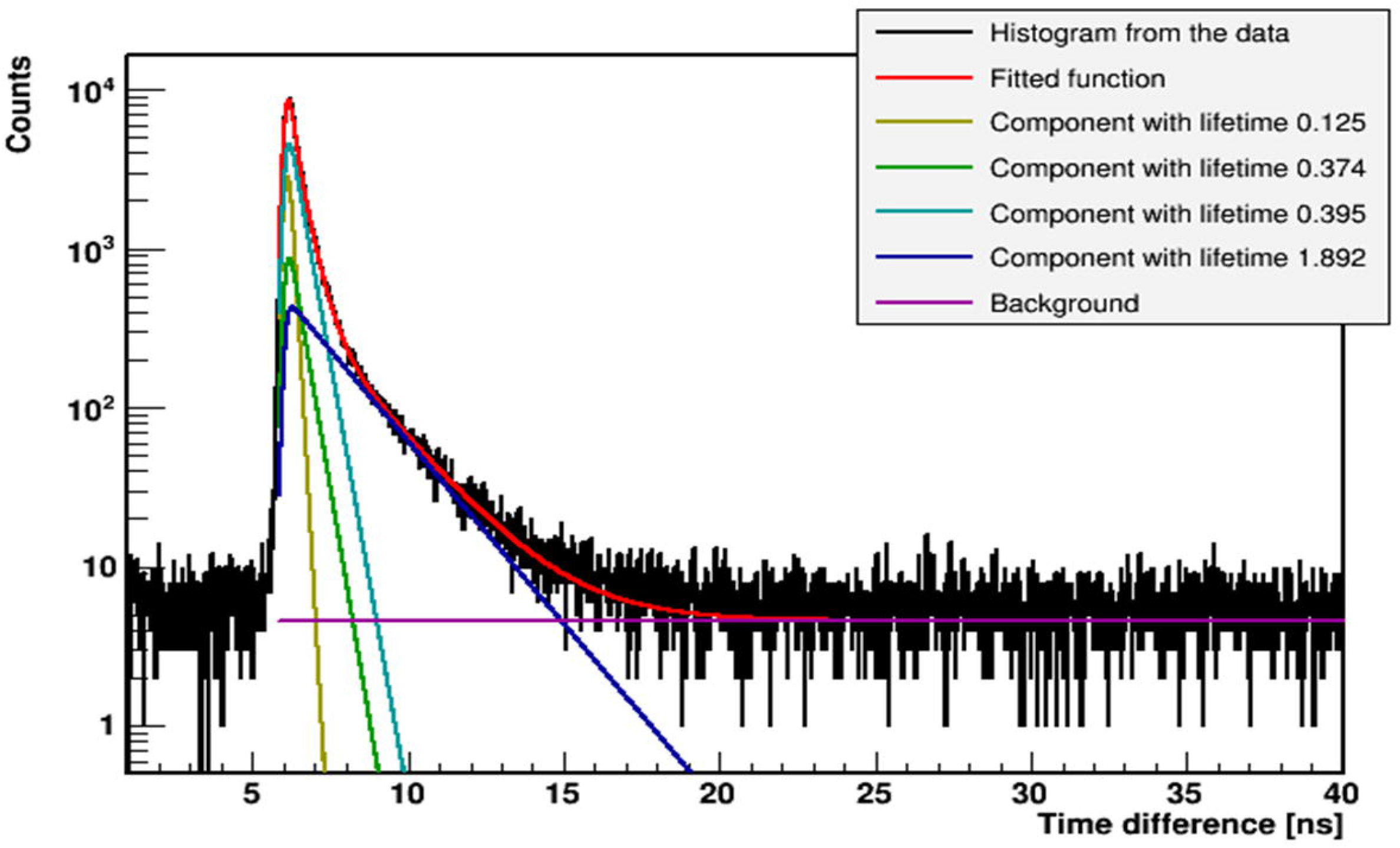
Exemplary experimental lifetime spectrum (black histogram) with superimposed histograms resulting from the fit of the sum of the exponential function convoluted with the detector resolution, performed by means of the PALS Avalanche program [24, 38-39]. First component (yellow line) shows the contribution from p-Ps (mean lifetime: 0.125 ns), second component (green line) originates from annihilations in source (Kapton foil) (0.374 ns), third component (light blue) shows free annihilation lifetime (0.395 ns), forth component (dark blue) illustrates the contribution from o-Ps. A sum of all contributions resulting from the fit is shown as a red curve.

Before each measurement, the system was calibrated, using an empty chamber including only the ^22^Na source. In order to perform the measurement, spheroids were transferred from the centrifuge tube to the chamber by using the scalper and ^22^ Na radioactive sources was placed between the samples. 3D spheroids were placed adjacent to the source, with no air between the source and spheroids. Then the chamber was put inside a holder that was connected to a heater to keep cells at 37°C. Finally, the holder was located between two BaF_2_ detectors, as indicated in (Fig 6).

The lifetime of positronium in spheroids with different ages, 4 and 8 days after seeding cells, has been evaluated based on the 10^6^ events collected for each studied case.

Positronium lifetime was extracted from each recorded spectrum by the fit of the sum of four exponential functions convoluted with the detector resolution function [38]. The exemplary spectrum with the result of the fit is shown in (Fig 7). The fitted components correspond to the decay of Para positronium (yellow curve), direct annihilation in the source/Kapton foil (green curve), and the annihilation of ortho-positronium (dark blue). The full experiment was repeated three times.

## Discussion

Recently positronium imaging was introduced [12, 16, 40] and a first *in vitro* positronium images has been demonstrated [17, 25] opening new possibilities for improvement of cancer diagnosis by using positronium as a tissue pathology biomarker [18].

The main aim of this study was to test the hypothesis that the positronium may serve as a novel biomarker for assessing the different cancer activity and biological properties in cancer cells and to perform first studies of positronium properties in the 3D cell spheroids.

3D spheroids were used as they can mimic the structure of real tumors under physiological conditions. The morphology and cell-cell interactions in 3D spheroids are completely different than in monolayer cell culture. 3D spheroids hold certain advantages over standard research methods, including low cost of use, high reproducibility, and timesaving, and reduces the need for laboratory animal models. The unique properties of 3D tumor spheroids make them invaluable for biological experiments and drug tests in a variety of experimental studies focused on chemotherapy and radiotherapy [41, 42].

The cell cycle time in 2D cells and 3D spheroids is significantly different because cells in monolayer cell culture have the homogenous biological condition and most of the cells are in the same cell cycle and cells have sufficient accessibility to nutrients and oxygen while in 3D cell culture with a multilayer structure, cells are in different cell cycle and accessibility to necessary nutrients for survival depend on their distance from the surface.

The rate of cell growth in WM115 spheroids is smaller (the doubling time is longer) than in WM266-4 spheroids, and WM115 cells spend more time in their cell cycle than WM266-4 cells. During the cell cycle, cells in the G1 phase are growing up the end of G2 phase when the mitosis and division starts after passing the checkpoints [43]. Spheroids have a necrotic core, a quiescent layer that cells are arrested in G1 phase of the cycle, and proliferating rim at the outermost part of spheroids that cells can be in S/G2/M [44].

The prior positronium lifetime research has been mostly done on the different type of tissues and monolayer cell culture such as skin cancer and colorectal cancer cells [22, 40, 45-49]. In this novel research, the lifetime of positronium has been evaluated in 3D spheroids rather than 2D cell culture or tissue. Although lots of research have been performed on positronium in materials and biological samples, the research using 3D cell aggregates is limited to the determination of o-Ps lifetime in 3D colorectal cancer cell aggregates with a mixture of cells and collagen [19]. We have performed also additional measurements to compare the o-Ps lifetime in 2D and 3D cell cultures. The obtained results, (Fig 8), demonstrate that positronium lifetime is smaller in 3D cell culture than in 2D. This graph shows how the o-Ps lifetime is different between 2D and 3D cell cultures which it backs to their diversity in biological properties and metabolism.

**Figure 8.**
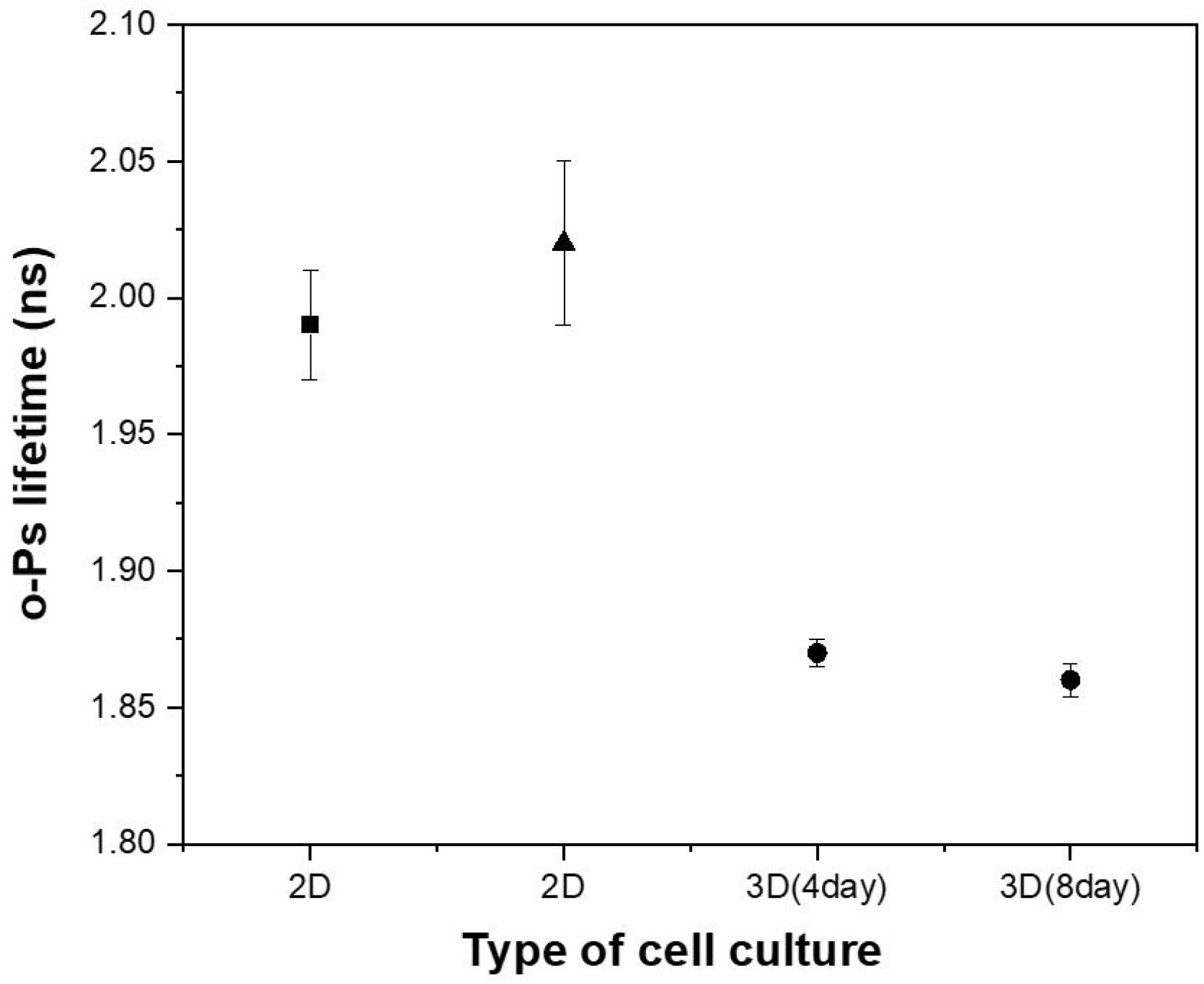
The lifetime of o-Ps in WM266-4 2D cell culture (black square) according to [50], the lifetime of o-Ps in WM266-4 2D cell culture (black triangle), and the lifetime of o-Ps in WM266-4 3D cell culture (black dots) show the o-Ps lifetime in two different ages of spheroids, 4 and 8 days after cell seeding.

This study has been conducted to evaluate the lifetime distribution of ortho-positronium in WM266-4 and WM115 melanoma cell spheroids characterized by different degree of malignancy. The measurements were performed at two different times (4 and 8 days) after seeding. During the time, both spheroid cell lines showed a reduction in o-Ps lifetime which backs to the gradual cell proliferation in spheroids. More malignant WM266-4 spheroids show smaller o-Ps lifetime in comparison to WM115 cell line spheroids.

In summary, the results show that the lifetime of o-Ps is useful parameter to differentiate between cancer cells with different biological characteristics. The higher the degree of malignancy and the rate of proliferation of neoplastic cells the shorter the lifetime of ortho-positronium. This research paves a way for development of positronium biomarker using cell spheroids model.

## Acknowledgements

This work was supported by the Foundation for Polish Science (FNP) through the TEAM POIR.04.04.00-00-4204/17 program, by the National Science Centre of Poland through grants no. 2019/33/B/NZ3/01004, 2021/42/A/ST2/00423, and the Jagiellonian University via project CRP/0641.221.2020 and DSC grant, number N17/MNS/000023 funded by Jagiellonian University which was awarded to Mrs. Hanieh Karimi for her PhD research.

## References

1. Stepien E, Karimi H, Leszczynski B and Szczepanek M. Melanoma spheroids as a model for cancer imaging study. Acta Phys. Pol. B. 2020;51(1):159–164.

2. Kapałczyńska M, Kolenda T. 2D and 3D cell cultures–a comparison of different types of cancer cell cultures. Arch Med Sci. 2018;14(4): 910–919.

3. Upreti M, Jamshidi-Parsian A, Koonce N, Weber J, Sharma S, et al. Tumor-endothelial cell three-dimensional spheroids: new aspects to enhance radiation and drug therapeutics. Transl. Oncol. 2011;4(6): 365–376.

4. Kunz-Schughart L.A, Kreutz M, Knuechel R. Multicellular spheroids: A three-dimensional in vitro culture system to study tumour biology. Int. J. Exp. Pathol. 1998;79(1):1–23.

5. Rastrelli M, Tropea S, Rossi C.R, and Alaibac M. Melanoma: epidemiology, risk factors, pathogenesis, diagnosis and classification. In vivo. 2014;28(6):1005–1011.

6. Voss R, Woods T, Cromwell K, Nelson K, Cormier J. Improving outcomes in patients with melanoma: strategies to ensure an early diagnosis. Patient Relat. 2015; 6:229–242.

7. Wróbel S, Przybyło M, and Stepien E. The clinical trial landscape for melanoma therapies. J. Clin. Med. 2019 8(3):368–382.

8. Karimkhani C, Green A, Nijsten T, Weinstock M, et al. The global burden of melanoma: results from the global burden of disease study 2015. Br. J. Dermatol. 2017 177(1):134–140.

9. Vörsmann H, Groeber F, Walles H, Busch S, Beissert S, et al. Development of a human three-dimensional organotypic skin-melanoma spheroid model for in vitro drug testing. Cell Death Dis. 2013; 4(7): e719.

10. Meghnani V, Vetter S, Leclerc E. RAGE overexpression confers a metastatic phenotype to the WM115 human primary melanoma cell line. Biochim Biophys Acta Mol Basis Dis. 2014;1842(7):1017–1027.

11. Moskal P, Jasińska B, Stępień E, Bass S. Positronium in medicine and biology. Nat. Rev. Phys. 2019;1: 527–529.

12. Moskal P, Kisielewska D, Curceanu C, Czerwiński E, Dulski K, et al. Feasibility study of the positronium imaging with the J-PET tomograph. Phys. Med. Biol. 2019; 64(5):055017.

13. Jasinska B, Gorgol M, Wiertel M, Zaleski R, Alfs D, et al. Determination of the 3 γ fraction from positron annihilation in mesoporous materials for symmetry violation experiment with J-PET scanner. Acta Phys. Pol. B. 2016;47(2):453–60.

14. Stepanov P.S, Selim F.A, Stepanov S.V, Bokov A.V, Ilyukhina O.V, et al. Interaction of positronium with dissolved oxygen in liquids. Phys. Chem. Chem. Phys. 2020;22(9):5123–31.

15. Shibuya K, Saito H, Nishikido F, Takahashi M, Yamaya T. Oxygen sensing ability of positronium atom for tumor hypoxia imaging. Commun. Phys. 2020 ;3(173):1–8.

16. Moskal P, Kisielewska D, Shopa RY, Bura Z, Chhokar J, et al. Performance assessment of the 2 γ positronium imaging with the total-body PET scanners. EJNMMI Phys. 2020;7(1):1–6.

17. Moskal P, Dulski K, Chug N, Curceanu C, Czarwinski E, et al., Positronium imaging with the novel multiphoton PET scanner. Sci. Adv. 2021; 7: eabh4394.

18. Moskal P, Stępień E. Positronium as a biomarker of hypoxia. Bio-Algoritms and Med-Systems. 2021;17(4):314–319.

19. Axpe E, Lopez-Euba T, Castellanos-Rubio A, Merida D, Garcia J.A, et al. Detection of atomic scale changes in the free volume void size of three-dimensional colorectal cancer cell culture using positron annihilation lifetime spectroscopy. PLoS One 2014;9(1): e83838.

20. Jean Y, Ache H.J. Positronium Reactions in Micellar Systems1. J. Am. Chem. Soc. 1977; 99(23): 7504–7509.

21. Liu G, Chen H, Chakka L, Gadzia J.E, Jean Y.C. Applications of positron annihilation to dermatology and skin cancer. Phys. Status Solidi C. 2007;4(10): 3912–3915.

22. Jean Y.C, Li Y, Liu G, Chen H, Zhang J, Gadzia J.E. Applications of slow positrons to cancer research: Search for selectivity of positron annihilation to skin cancer. Appl. Surf.Sci. 2006;252(9):3166–3171.

23. Jasińska B, Zgardzińska B, Chołubek G, Gorgol M, Wiktor K, et al. Human tissues investigation using PALS technique. Acta Phys. Pol. B 2017; 48:1737–1748.

24. Moskal P, Kubicz E, Grudzien G, Czerwinski E, Dulski K, et al. Developing a Novel Positronium Biomarker for Cardiac Myxoma Imaging. bioRxiv 2021.

25. Moskal P, Gajos A, Mohammed M, Chhokar J, Chug N, et al. Testing CPT symmetry in ortho-positronium decays with positronium annihilation tomography. Nat. Commun. 2021; 12(1):1–9.

26. Matulewicz T. Radioactive nuclei for β+ γ PET and theranostics: selected candidates. Bio-Algoritms and Med-Systems. 2021;17(4):235–9.

27. Choiński J, Łyczko M. Prospects for the production of radioisotopes and radiobioconjugates for theranostics. Bio-Algoritms and Med-Systems. 2021;17(4):241–57.

28. Karp J.S, Viswanath V, Geagan M.J, Muehllehner G, Pantel AR, et al. PennPET explorer: design and preliminary performance of a whole-body imager. J. Nucl. Med.. 2020;61(1):136–43.

29. Moskal P, Stępień E.Ł. Prospects and clinical perspectives of total-body PET imaging using plastic scintillators. PET Clin. 2020 ;15(4):439–52.

30. Badawi R.D, Shi H, Hu P, Chen S, Xu T, et al. First human imaging studies with the EXPLORER total-body PET scanner. J. Nucl. Med. 2019;60(3):299–303.

31. Vera D.B, Schulte B, Henrich T, Flavell R, Seo Y, et al. First-in-human total-body PET imaging of HIV with 89Zr-VRC01 on the EXPLORER. J. Nucl. Med. 2020; 61:545.

32. Vandenberghe S, Moskal P, Karp JS. State of the art in total body PET. EJNMMI phys. 2020; 7:1–33.

33. Sakai Y, Nakazawa K. Technique for the control of spheroid diameter using microfabricated chips. Acta Biomater. 2007; 3(6):1033–1040.

34. Song Y.S, Park C.M, Park S.J, Lee S.M, Jeon Y.K, Goo J.M. Volume and mass doubling times of persistent pulmonary subsolid nodules detected in patients without known malignancy. Radiology 2014;273(1):276–84.

35. Mehrara E, Forssell-Aronsson E, Ahlman H, Bernhardt P. Specific growth rate versus doubling time for quantitative characterization of tumor growth rate. Cancer res. 2007;67(8):3970–5.

36. Mordecai Schwartz. A biomathematical approach to clinical tumor growth. Cancer, 14(6):1272–1294, 1961.

37. Karimi H, Leszczyński B, Kołodziej T, Kubicz E, Przybyło M, Stępień E. X-ray microtomography as a new approach for imaging and analysis of tumor spheroids. Micron. 2020;137(10): 102917.

38. Dulski K. PALS avalanche: a new PAL spectra analysis software. Acta Phys. Pol. A. 2020;137(2):167–170.

39. Dulski K, Bass SD, Chhokar J, Chug N, Curceanu C, et al. The J-PET detector—a tool for precision studies of ortho-positronium decays. Nucl. Instrum. Methods. 2021; 1008:165452.

40. Moskal P. Positronium imaging. In 2019 IEEE Nucl. Sci. Symp. Med. Imaging Conf. Proc. (NSS/MIC) 2019: 1–3.

41. Costa E, Moreira A, Melo-Diogo D, Gasper V, Carvelho M, Correia I. 3D tumor spheroids: an overview on the tools and techniques used for their analysis. Biotech. Adv. 2016;34(8):1427–1441.

42. Pampaloni F, Ansari N, and Stelzer E. High resolution deep imaging of live cellular spheroids with light-sheet-based fluorescence microscopy. Cell Tissue Res. 2013; 352(1):161–177.

43. Ginzberg MB, Kafri R, Kirschner M. On being the right (cell) size. Cell Biol. 2015;348(6236):771–780.

44. Beaumont K.A, Anfosso A, Ahmed F, Weninger W, Haass N.K. Imaging-and flow cytometry-based analysis of cell position and the cell cycle in 3D melanoma spheroids. J. Vis. Exp. JOVE.2015(106): e53486.

45. Sane P, Tuomisto F, Wiedmer S, Nyman T, Vattulainen I, Holopanainen J. Temperature-induced structural transition in-situ in porcine lens—changes observed in void size distribution. Biochim Biophys Acta Biomembr 2010;1798(5): 958–965.

46. Elias M, AL-Mashhadani A, AL-Shiebani Z. Temperature Dependence of Microstructure of Biological Tissues Probed Via the Positronium Method. Pure Sci. 2001; 28(2): 240–244.

47. Chandrashekara M.N, Ranganathaiah C. Free volume size distribution in some natural polymers. AIP Conf Proc. 2011;1349: 228–229.

48. Chandrashekara M.N, Ranganathaiah C. Diffusion of permanent liquid dye molecules in human hair investigated by positron lifetime spectroscopy. Colloids Surf. B. 2009; 69(1):129–134.

49. Torisawa Y, Takagi A, Nashimoto Y, Yasukawa T, Shiku H, Matsue T. A multicellular spheroid array to realize spheroid formation, culture, and viability assay on a chip. Biomaterials 2007; 28(3):559–566.

50. Ewelina Kubicz. Biomedical applications of Positron Annihilation Lifetime Spectroscopy: nano structural characterization of normal and cancer cells and tissues. PhD thesis, Jagiellonian University, 2020.

